# Unsupervised inference of protein fitness landscape from deep mutational scan

**DOI:** 10.1101/2020.03.18.996595

**Authors:** Jorge Fernandez-de-Cossio-Diaz, Guido Uguzzoni, Andrea Pagnani

## Abstract

The recent technological advances underlying the screening of large combinatorial libraries in high-throughput mutational scans, deepen our understanding of adaptive protein evolution and boost its applications in protein design. Nevertheless, the large number of possible genotypes requires suitable computational methods for data analysis, the prediction of mutational effects and the generation of optimized sequences. We describe a computational method that, trained on sequencing samples from multiple rounds of a screening experiment, provides a model of the genotype-fitness relationship. We tested the method on five large-scale mutational scans, yielding accurate predictions of the mutational effects on fitness. The inferred fitness landscape is robust to experimental and sampling noise and exhibits high generalization power in terms of broader sequence space exploration and higher fitness variant predictions. We investigate the role of epistasis and show that the inferred model provides structural information about the 3D contacts in the molecular fold.

## I. MAIN

The continuous interplay between selection and variation is at the basis of Darwinian evolution. Recent advances in experimental techniques allow for a quantitative assessment of evolutionary trajectories at the molecular level [1]. From this point of view, the improvement in the construction of combinatorial libraries of proteins or other biomolecules and the high-throughput technologies to characterize their phenotypes [2, 3], provides one of the more stringent testing grounds for studying the genotype-phenotype evolutionary relation under an externally controlled selective pressure. Besides its evident theoretical appeal, this line of research also has a more practical interest: in Directed Evolution experiments, combinatorial libraries of sequences are routinely screened to select molecules with specific biochemical properties such as binding affinity towards a target (*e.g.* antibodies) [4], catalytic features (*e.g.* enzymes) [5–7], etc.

The last decades have seen a tremendous boost in the availability of reliable high-throughput selection systems, such as: genetic [8], display systems (e.g. phage, SNAP-tag, mRNA, etc) [9], cytofluorimetry (e.g. FACS) [10], and micro-droplet techniques [11]. Still, a fundamental limitation is the number of variants that can be screened compared to the size of the sequence space of possible mutants. For example, a hundred residue protein has up to 20^100^ ~ 10^130^ possible variants, while the actual massive parallel assay libraries are typically able to handle ranges of variability within 10^8^–10^12^, which can be fed to a single high throughput screening pass.

Thanks to advances in sequencing technologies (especially in terms of reduced cost per read), machine learning methods for the inference of sequence-phenotype associations are starting to show their full potential. In particular, several massively parallel assays, known as deep mutational scanning (DMS) [2, 3, 12] are becoming available where typically a large-scale library of protein variants undergo repetitive cycles of selection for a function or an activity. The library is retrieved each round and the counts of each variant are determined by high-throughput sequencing. Such an increasing amount of sequence data, demands new algorithms to produce accurate statistical models of genotype-fitness associations.

All the computational methods developed so far that make use of deep mutational scanning sequencing data to learn a genotype-fitness map utilize a supervised approach: a proxy of the fitness of the mutants tested in the experiment is computed from the sequencing reads and a machine learning method solves the regression problem [13–18] (with the only remarkable exception of [19] that we discuss later). Here, we propose a novel method that shifts the learning approach from a supervised to an unsupervised framework.

We learn a model that describes accurately (via maximum likelihood) the full-time series (successive rounds) of sequencing reads observed in the experiment.This strategy has the advantage of not reducing the information contents defining a function of the fitness that is often affected by sampling noise, but on the contrary, it uses the full information in the screening experimental data: the sequencing reads time series.

The method consists of a probabilistic modeling of the three phases of each experiment cycle: i) selection, ii) amplification, and iii) sequencing (see Methods). In brief, what we observe are the reads coming from the sequencing, *i.e.*, a sample of the library at a specific time step. The other phases are described in terms of latent variables referring to the number of amplified and selected mutants. The probability that a mutant is selected (*e.g.* by physical binding to the target) depends on the specific mutant sequence composition. On the other hand, we assume that the probability of a mutant to be amplified depends only on the fraction of mutants present after selection (ignoring possible sources of amplification selection such as codon bias that could however be taken into account in our framework using appropriate priors). We take into account both additive contributions from the individual residues and epistatic contributions in the form of pairwise interactions, although more complex multi-residue interaction schemes could be introduced.

This probabilistic description allows us to define an overall likelihood to observe a time series of reads in an experiment given the parameters involved in the *energetic* contribution to the selection, i.e. the genotype-phenotype map. Optimizing the parameters to maximize the likelihood allows us to obtain an effective model of the fitness landscape.

The method has the twofold aim of: (i) providing an accurate statistical description of the time series (in terms of panning rounds) evolution of the differential composition of the combinatorial library, (ii) predicting individual sequences, or rationally designed libraries of increased biophysical activity towards the sought target, that in particular, can be used in the recently proposed machine-learning-guided directed evolution for protein engineering [18].

The method we propose gets inspiration from the Direct Coupling Analysis (DCA) methods developed to describe statistical coevolutionary patterns of homologous sequences [20]. This successful field has provided fundamental tools commonly applied in structure prediction pipelines [21] and more recently to provide mutational effect predictions [22–27]. Other approaches apply different machine learning schemes [28] on the same framework. The main difference between these unsupervised methods and the present work lies in the input data. The DCA approach learns a statistical description of a multiple sequence alignment of the protein family sequences, treating it as if it were an equilibrium sample drawn from a Potts model, while we deal with an out-of-equilibrium time series of screening experiments reads. Moreover, the broadness of the sequence space covered by the input data is different, with a protein family typically reaching an average Hamming distances of around 70% which results from the outcome of millions of years of Darwinian evolution, while in the DMS typical combinatorial libraries have at most 4-5 mutations away of the wild types in a few rounds of selection.

This opens several questions on the relevance of the modeling. Notably on the role of epistatic interactions and the extent of applicability of the method as a generative model. The application of DCA methods has provided evidence on the importance of epistatic effects in shaping the homologs distribution over sequence space. Moreover, on the modeling side, it has shown the effectiveness of pairwise models [29, 30] to capture the epistatic contribution and provided useful three-dimensional structural predictions (residue contacts) [16, 17].

In the Results section, we investigate whether the same applies to the output of the experimental assay, first and foremost the reliability of the inferred epistatic interactions and the effectiveness to generate optimized sequences with respect to the selection process. Our findings are corroborated by the capacity to predict structural properties (e.g. residue contacts) from the inferred epistatic interaction, as similarly found in [16, 17].

## II. RESULTS

First we assess the accuracy of the inference method to learn the genotype to fitness map by testing on five Deep Mutational Scanning (DMS) datasets, briefly described in the Data section. Also, we investigate the generalization power of the inferred fitness landscapes and the promising potential to generate sequences of high fitness. Finally, we examine the epistatic interactions learned, comparing them to non-specific epistasis due to a global non-linear genotype to fitness map [14], and analyzing the relevance of the epistatic terms to predict contacts between residues in the three-dimensional molecular structure.

In a typical screening experiment, the selectivity [13] is a measure of the fitness of a protein mutant computed from the sequencing samples of the population. In its simpler form, it is the ratio of the sequence counts at two consecutive rounds. Slightly different definitions of selectivity are present in the literature [13], aiming to reduce the impact of experimental noise on the computed mutants fitness.

There are several sources of noise that affect the reproducibility of a deep mutational scanning experiment. Sequences that are present in low numbers are more susceptible to statistical fluctuations. This can be due to the uneven initial library composition that can change between different realizations of the same experiment. In addition, the attempts to cover a large sequence space can generate low replicates per mutants, since the availability of particles to carry the mutants is limited (*e.g.*, in practice no more than about 10^13^ phages can be manipulated). Moreover, the sequenced mutants represent a very small sub-sample of the total diversity of variants in the experiment. Therefore the reads statistics might not reflect fairly the underlying variant abundances. The magnitude of this sampling error depends on the *mutants coverage* that we define as the ratio of the total number of reads over the number of unique mutants, i.e. the number of reads per variant in a hypothetical uniform distribution case. In Table (I), we list the mutants coverage for each used data set. The sampling noise affects both the trained model and the selectivity measure but, interestingly, as we see later has a more prominent effect on the latter.

**TABLE I.** Different deep mutational scanning datasets used in the paper to evaluate the performance of the model.

| Reference | Protein | Target | Length mutated part | Selection rounds | Sequenced time points | Unique mutants | Average distance from wild type | Mutant coverage |
| --- | --- | --- | --- | --- | --- | --- | --- | --- |
| Olson et al. (2014) | GB1 | IgG-Fc | 55 | 1 | 2 | 536833 | 2 | 588 |
| Wu et al. 2016 | GB1 | IgG-Fc | 4 | 2 | 2 | 157161 | 3.8 | 422 |
| Fowler et al. (2010) | WW domain | peptide | 25 | 6 | 3 | 572076 | 3.4 | 14 |
| Araya et al. (2012) | WW domain | peptide | 34 | 3 | 4 | 940730 | 4.3 | 11 |
| Boyer et al. 2016 | Ab IgH | PVP, DNA | 4 | 3 | 3 | 28195 | 3.5 | 4 |

### Validation of mutants fitness predictions

To validate the inferred genotype-phenotype map we do not have access to direct high-throughput measures of the binding energy with the target. Nevertheless, we can assess the reconstructed fitness landscape comparing the predicted binding energy with out-of-sample sequence selectivities. To do so, we perform a leave-one-out 5 fold cross-validation, *i.e*, masking a fifth of the mutants in the library from the learning data, and testing on the remaining ones. To mitigate the effect of the sampling noise on the selectivity measure, we filter out sequences with high selectivity error (see Methods for details) from the test set. In figure (1), we show the correlation of the predicted binding energy and the log selectivity for each examined data set. In all experiments, we obtain an excellent agreement between the out-of-sample model prediction and the selectivity measure based on read counts (i.e. a proxy of the binding energy). The correlation between the two values steadily increases as we filter out more noisy sequences.

**FIG. 1.**
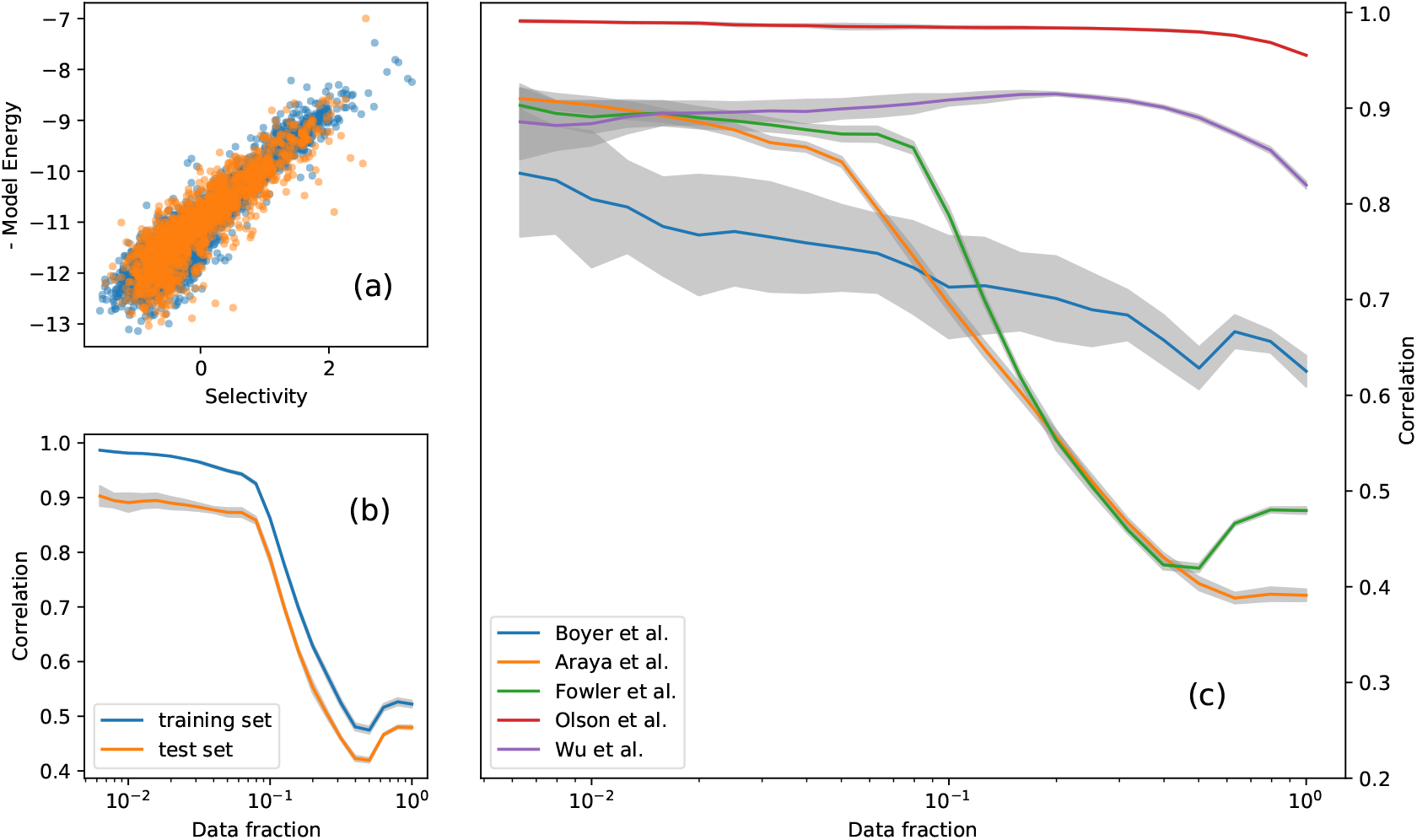
Overall model performances. Correlation of the predicted binding energies *E* and the log-selectivity *θ* computed from the sequencing reads. **a)** Scatter plot of *E* vs *θ* for the in-sample (blue dots) and out-of-sample (orange dots) sequences in the Araya et al. dataset. The Pearson correlation is *ρ* = 0.81, after filtering out the noisier data (fraction of used data *f* = 0.05). **b)** Pearson correlation coefficient between *E* and *θ* for different filtering thresholds on sequence errors in the Araya et al. dataset. On the x-axis, the fraction of the sequences used to compute the correlation, a lower fraction of sequences account for a more severe filter on noisy sequences and provide higher correlations. The comparison of the in-sample and out-of-sample sets (4/5 and 1/5 of the mutants respectively) shows a minor overfitting bias. **c)** Comparison of the Pearson correlation curves (same of panel b) for the five datasets. Notice that the higher the mutants coverage of the dataset (Olson et al., Wu et al.) higher the correlation reached, see table (I)

As previously pointed out, in contrast to other approaches that fit sequence selectivities, we train a model directly on the sequencing reads, maximizing the model likelihood of the full set of read counts from the experiment and obtaining a statistical description of the differential composition of the combinatorial library across rounds. This allows us to obtain an estimate of the binding probability more reliable and robust against the experimental noise. To prove this statement, we create a decimated training set by selecting in each round of the experiment a random subset of the reads, and for each training-set realization we learn the model parameters. We perform the test on the Olson et al. dataset, which has a high mutants coverage (~ 500).

The results are shown in figure (2): from panels (a-b) we clearly see that the reliability of the selectivity decays faster as the decimation ratio increases, compared with that of the model which provides accurate predictions also in the highly under-sampled regime. In other terms, the selectivity of a sequence derived from an under-sampled dataset is a worst statistical predictor of the full-dataset selectivity compared to our model predictor. Even if we use the correction strategy outlined by [13], the selectivity measure is severely impacted by sampling noise, whereas the predicted fitness landscape inferred by our model is more robust to under-sampling. Finally, the measures of selectivity relies upon the presence of multiple reads of the same mutant across the rounds, while our approach seems to be less impeded by this limitation.

**FIG. 2.**
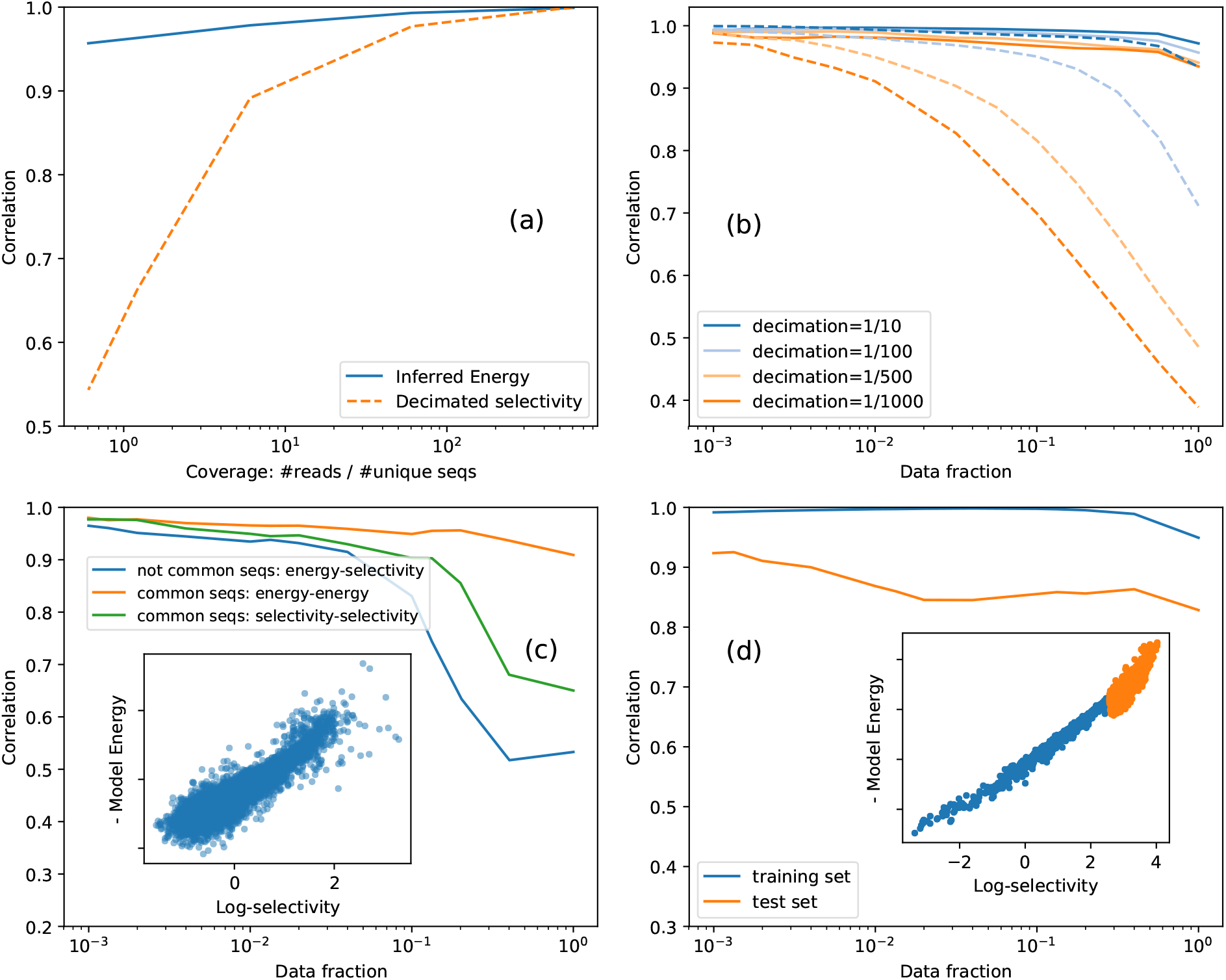
Model robustness and generalization power. In panels **(a-b)** is shown the robustness of the model inference with respect to sampling noise. We reduce the mutants coverage decimating randomly the reads in the Olson et al. dataset, obtaining datasets with different numbers of total reads. We use them to infer the binding energy and to compute the selectivity after decimation and we compare them with the full dataset selectivity. In panel **(a)** we show the Pearson correlation coefficient between the full dataset selectivity and the predicted energy of the model and the decimated selectivity on a test set, as a function of the coverage of the decimated datasets. In panel **(b)** we show the correlations of the two measures with the full dataset selectivity for different thresholds of filtering of mutants errors on the test set. While the Pearson correlation between the decimated dataset selectivity and the original one reduces drastically upon increasing the decimation rate, the predicted binding energy maintains always a high correlation with the test-set selectivity. **c)** Correlation of predicted energies *E* and empirical selectivity *θ*, when the model is trained on one dataset (Araya et al.) and tested on the outcome of a different experiment(Fowler et al.). On the *x*-axis, the fraction of the sequences used to compute the correlation after filtering on error. The blue curve refers to the model trained on sequences that are not common to the two datasets. In the inset a scatter plot of *E* and empirical log-selectivity for a particular choice of filter threshold (data fraction *f* = 0.05, correlation *ρ* = 0.91). The yellow curve corresponds to the correlation between inferred energies of the sequences common to the two datasets while the green one refers to the correlation between selectivity compute on the same sequences in the two datasets, interestingly is lower than the previous correlations suggesting that energy is a more reproducible quantity than the empirical selectivity itself. **d)** Capacity of the model to predict best binders. Correlation of predicted energies *E* and empirical selectivity *θ* trained on low selectivity mutants and test on the top selectivity ones. The high correlation in the test set and the capacity to rank properly the unseen best binders suggest the promising application of the method as a generative model.

### Generalization extent of the inferred fitness landscape

The fitness landscape refers to the selection process of a specific phenotype of a protein. We investigate to which extent the model is able to extract information about general features of the landscape that can be used to provide reliable predictions on different experiments, different energy spectra or different sequence space regions with respect to the training ones.

An intriguing question is to what extent different experimental settings can have an impact on the inferred parameters, reducing its generalization power, or whether we can use the learned map to predict new experimental outcomes.

Fowler et al. [31] and Araya et al. [32] published two experimental datasets, in which the hYAP64 WW domain is selected for binding against its cognate polyproline peptide ligand. In both cases, most variants in the library are on average two a.a. substitutions away from the wild-type sequence. Still, the initial libraries of the two experiments have only about 50% sequences in common, with the rest of the sequences being unique to each dataset. The model trained on one dataset (discarding the common sequences) provides accurate predictions of the empirical selectivities observed in the other experiment, as shown in panel (a) of figure 2. Interestingly when the common sequences are taken into account, the binding energies inferred from the two datasets show better correlations than the selectivities. This suggests that the inferred energy is more reproducible and robust to noise, and hence is a better estimator of the *true* fitness.

We repeated the same analysis training on the Olson et al. dataset and testing on the Wu et al. dataset, as they both used the IgG-binding domain of protein G (GB1) and performed the selection for binding to immunoglobulin G fragment crystallizable (IgG-Fc). In this case, we train on all the sequences at Hamming distance 1 and 2 from the wild-type (from the Olson et al. dataset) and we ask whether the model can make predictions of sequences three or four mutations away from the wild-type (from the Wu et al. dataset). The results in figure (4) of the SI show that the model is still able to predict the fitness landscape for more distant mutants (Pearson correlation of *ρ* = 0.67 for Hamming distance 3 and *ρ* = 0.55 for Hamming distance 4), although these predictions deteriorate as the distance to the sequences covered in the training set increases.

Can we learn from sequences in the low binding energy-band a predictive model of the high binding energy-band of the fitness landscape? This is a relevant question if we want to exploit the model to generate, for instance, better binders not originally present in the experiment. To gauge the performance of the model for this task, we use the sequences with low selectivity as training set, and the sequences with higher selectivity as the test set.

In contrast to the previous results where the out-of-sample sequences were extracted from the same distribution as the in-sample, in the present computation we selectively learn from low-medium binding energy sequences while we test on the top binders. As shown in panel (c), figure (2), the predictions are in excellent agreement with the data, showing the capacity to learn the fitness landscape of high fitness region very accurately.

### Epistasis

The study of the role of intragenic epistatic interactions in shaping the fitness landscape is a subject of intense research, different contrastive results have been largely debated in the scientific community [4, 14, 33–35]. Although on the evolutionary scale it is clear that the epistasis have an important role in shaping the sequence ensemble of protein family domains across homologs, on a more local scale, in the selection of local mutations around a wild type sequence for binding a target, is debated whether such effects are involved and to which amount. We investigated whether a model without epistasis, hence where the mutation effects are independent in each residue and provide additive contributions to fitness, can reach the same description accuracy of the experiments. See Methods for details on the independent site model and the pairwise epistatic one.

The five datasets considered in this study vary with respect to the broadness of sequence space sampled (how far from the wild type are the mutants in the library) and the length of the mutated part of the sequence, as summarized in table (I).

The two opposite limits are the Olson *et al.* dataset where the full length of the gb1 (55aa) is mutated only by a maximum of two amino acids (long sequence, limited broadness) and the Boyer *et al.* and Wu *et al.* datasets where only four amino acids are considered but the libraries cover a significant fraction of sequence space. The Araya *et al.* and Fowler *et al.* datasets lay in an intermediate regime.

Panels (a-c) of figure 3 show the comparison of the performance of the independent site model and the pairwise epistatic model: the broader the sequence space covered in the experiment, the more crucial becomes the inclusion of the epistatic interactions in the model for a proper description of the experimental outcome.

**FIG. 3.**
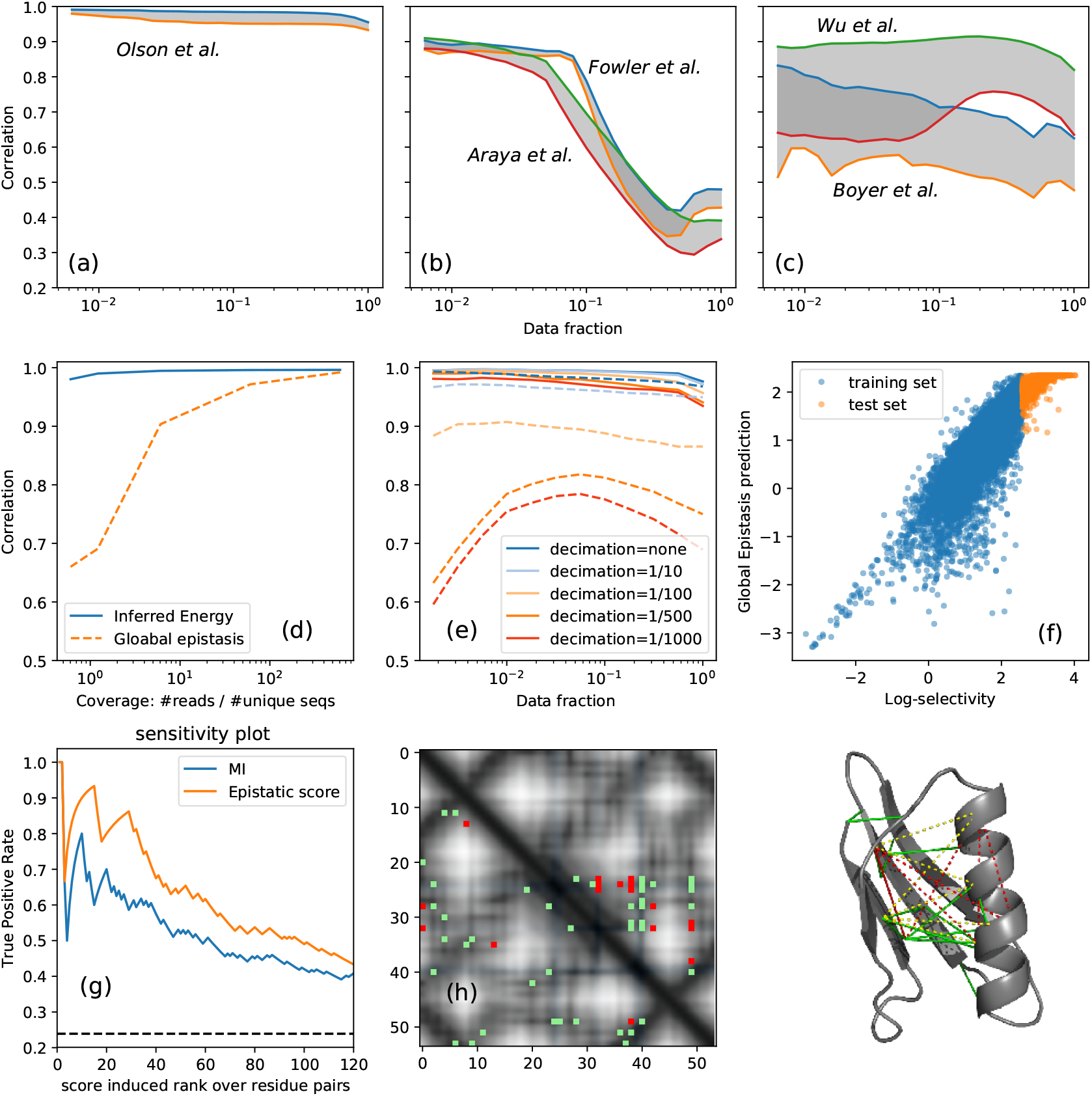
Epistatic Effects. The comparison of the independent site model and the epistatic model is displayed in panels **(a-c)**. The three panels refer to different characteristics in the broadness of the screened library and the length of the mutated part of the protein sequence. From left to right broadness increase and the number of the mutated residues reduce (see table I. For each dataset are shown the correlation of the epistatic model (upper line) and the independent site model (lower line) and is highlighted in gray the gap between the two. Broader the library, more distant mutants from the wild type are screened and more the epistatic effects become relevant. In panels **(d-f)** are shown the robustness and generalization analysis (same as depicted for the epistatic model in figure 2) of the global epistasis model [14]. The panels **(d-e)** display the reduction of the accuracy of GE model when lowering the mutants coverage. In panel (f), the low correlation between GE prediction and selectivity for high fitness mutants, depict the deficiency to generalize the prediction to lower binding energy spectra. The panels **(g-i)** show the structure contacts predictions for GB1 domain form the Olson et al. DMS experiment (tested on the crystal structure PDB id: 1fcc). Panel **(g)** : the ROC curves of the predicted contacts with the epistatic score computed from the inferred model and the weighted Mutual Information. Panel **(h)** : Contact map of the first predicted contacts. In gray-scale is displayed the distances between residue heavy atoms, in blue are highlighted the residues on the binding surface (less than 3Å to the Fc domain of human IgG). The green dots are the true positives (heavy atoms distance less than 8Å) and the reds are false positives. In the upper triangular part are shown the Mutual Information predictions while in the lower triangular part the Epistatic score. The MI predictions are strongly clustered around the binding surface while the model predictions cover the whole structure. Panel **(i)** The same predictions on the molecular structure of G protein in complex with the Fc domain of human IgG.

Recent papers have pointed out that epistatic interactions can arise spuriously from non-linearities in the genotype-phenotype map [14, 33].

Otwinowsky proposed a global epistatic model that infers the parameters of an independent site model together with the nonlinear shape of the map. This method provides a good prediction of the fitness in DMS experiments (See figure (5) in SI for the performance on all the datasets), considering also the lower number of parameters used. Nonetheless, performing the same analysis to test robustness and generalization highlighted in the previous paragraph, the global epistasis model appears to be sensitive to sampling error (panels (e-f) of figure 3) and fails to predict higher fitness mutants when trained on the low fitness ones (panel (f) of figure 3)

In recent studies, it has been demonstrated that the epistatic interactions, quantified from deep mutational scanning experiments, can be used to determine 3D contacts in the molecular structures [15–17]. This finding provided a strong support to the idea that the observed epistatic interaction does not come only from non-specific artifacts due to the non-linearity of the fitness map, but rather reflects the interplay of structural stability and functional binding of the selection process in the experiment.

We investigate whether and to which extent the proposed model provides contact predictions that can be used for three-dimensional structure modeling, on the GB1 domain of protein G using the Olson et al. DMS experiment. To test the predictions we use a crystal structure of the protein in complex with the Fc domain of human IgG (PDB id 1fcc). As a measure of the epistasis between two positions we use the average difference in binding energy between the sum of single mutations and the double mutants, similar to the score used by [36] (see the methods section for details). We identified the most epistatic pairs by sorting all of the pairs of positions by order of decreasing epistatic score.

In figure 3 panel (g) we show the ROC curve of the predictions compared to the weighted Mutual Information (a non-parametric measure defined in equation (S.10)). In panel (h) are shown the first 20 predictions and the true contact maps. We note that the MI predictions are strongly clustered around the binding surface, while the model contact predictions are distributed all over the structure, making them more useful for 3D structure modeling. In the supplementary material, we report the same structural analysis for the WW domain, using the dataset from Fowler *et al.* and Araya *et al.*.

## III. DISCUSSION

Despite advances in high-throughput screening and sequencing techniques, investigating genotype-phenotype relationships remains a substantial challenge due to the enormous size and large dimensionality of the space of possible genotypes. We propose a computational method to obtain a model of the genotype to fitness map, learned from the sequencing data of a deep mutational scanning experiment. The novelty of the method consists of an unsupervised approach that uses a probabilistic description of the full amplification-selection-sequencing phases of the experiment. One key element lies in the inclusion of pairwise epistatic interactions in the modeling of the specific mutant selection, nevertheless we remark that in the same framework other modeling schemes are possible.

To investigate the properties of the inferred fitness landscape, we used five DMS experiments related to two well-studied proteins, the WW domain part of the YAP65 protein, the Gb1 domain of the IgG-binding protein, and the variable part of a human antibody, all selected for binding to cognate ligands. These experiments differ in several technical characteristics, such as library generation and expression, length of the mutated part of the protein, broadness of the initial library and sequencing coverage per mutants.

First, we performed a cross-validation test obtaining accurate selectivity predictions for all the five datasets. Second, we investigated the generalization power of the model. We learned on one experiment and predicted correctly the outcome of a second one with same wild type protein and binding target, obtaining a better experimental reproducibility than from the mutant selectivities themselves. Remarkably the model shows the capacity to predict lower binding energy spectra, we masked the higher fitness mutants from the training and we recover the correct ranking and fitness of the best binders. Moreover, we noticed that the predicted fitness landscape is more robust to experimental noise than the selectivity measures (the fitted quantity in the supervised approach). To demonstrate this we performed a decimation of sequencing reads and assess the detrimental in the predictions.

Finally, we investigated the reliability of the epistatic interactions in the model. Our results show that when increasing the library’s sequence diversity, epistatic interactions become more important to obtain a good fit to the experiments. In addition, we can extract from the inferred epistatic interactions, structural information of the 3D contact proximity.

Recently it has been pointed out that non-linearities in the genotype to fitness map can produce spurious epistatic effects, namely non-specific epistasis or global epistasis [14, 33]. This suggests a limited magnitude of specific epistatic effects in shaping the fitness landscape in local screening assays. We compared the two hypothesis and our analysis suggest that, while the spurious global epistasis could have a prominent role where the experiment is selecting *complex* phenotypes (among others: cell growth rate [37, 38] or a proxy of expression levels [39]), in the set of experiments we have analyzed, where the selection is upon the binding affinity to a target molecule, the specific epistatic effects account for real genetic interactions.

All these findings suggest that the presented unsupervised approach has the promising application as a generative model to identify novel high-fitness variants and can be included in a machine-learning-assisted Directed Evolution framework where the computational part are included in the cycle to design the combinatorial libraries to be screened [40].

## IV. METHODS

### Inference

We consider a set of experimental rounds of selection *t* ∈ 0, 1, …, *T*, with *t* = 0 referring to the initial combinatorial library. At round *t*, there are 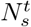 phages displaying sequence *s*. The number of phages that will be selected for the next round can be taken as binomially distributed with mean 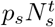, where *p*_*s*_ is the selectivity of sequence *s*. Since a large quantity of phages is present initially, we can approximate this as the deterministic selection of a fraction 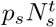 for each sequence. In the next step, the selected pool is thus amplified to reach the initial population size. Since a small quantity of carriers is selected, this step is modeled as a stochastic multinomial distribution,

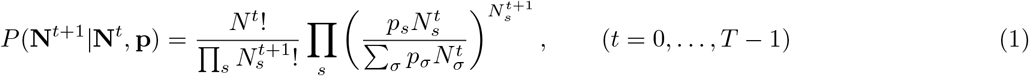

where 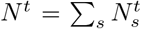 is the total number of particles carrying sequence *s* at round *t*, and a bold symbol such as **N**^*t*^ denotes the set of all 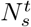 for all sequences. The full experiment consists of iterating these two steps.

Finally, at selected rounds, a sample of the amplified population is sequenced. In the limit of large enough sample size, we assume that the read counts are approximately proportional to the frequencies of sequences in the population (see Appendix for details).

Under these assumptions, it follows that the likelihood of the time-series {**N**^0^, **N**^1^, …, **N**^*T*^} is given by the product of (1) from *t* = 0 to *t* = *T* − 1.

#### Genotype to fitness map

The selection probabilities can be modeled by a two-state thermodynamic model (bound or unbound), 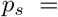 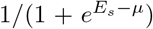, where *E*_*s*_ is the binding energy of sequence *s* and *μ* is the chemical potential, which depends on the concentration of binding target presented in the experiment [41].

As a function of sequence, *E*_*s*_ is a genotype-to-phenotype mapping that assigns a biophysical parameter (binding energy) to each sequence. We assume that *E*_*s*_ decomposes into additive contributions from individual a.a. species in the sequence, plus epistatic contributions from interacting pairs of letters:

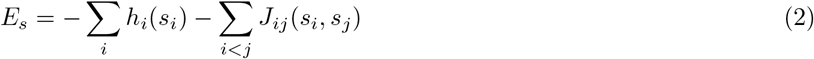

The problem becomes that of inferring the parameters *h*_*i*_(*a*), *J*_*ij*_(*a, b*) by maximizing the likelihood (1) over all rounds. In addition a regularization (e.g. an *ℓ*_2_-norm) term can be included to prevent over-fitting.

#### Rare binding approximation

In typical experiments, the fraction of selected phages is very small, implying that *p*_*s*_ ≪ 1 for most sequences. This suggests a rare binding approximation, 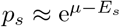. Under this approximation, the log-likelihood simplifies to:

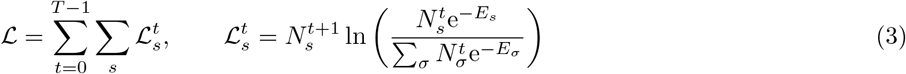

In this limit, the log-likelihood does not depend on *μ* anymore. It also makes *ℒ* a concave function of the energies *E*_*s*_, and hence of the fields *h*_*i*_(*a*), *J*_*ij*_(*a, b*) that we intend to learn. Since our algorithm consists in finding the maximum of (3) with respect to these parameters, concavity guarantees that the solution is unique and that it can be found efficiently with numerical optimization routines. In our implementation we found that the L-BFGS algorithm performed well.

#### Independent site model

Due to the rare binding approximation, when in the energy terms are considered only the *h* parameters contribution, each residue contributes independently and there are no epistatic effects as the amino acid changes impact additively to the fitness, 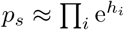.

#### Empirical selectivity

To compute empirical selectivities, we performed a linear regression of parameters *θ*_*s*_, *α*^*t*^, in a model of the form:

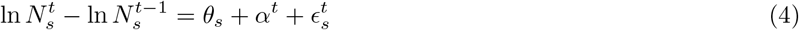

where *θ*_*s*_ is the log-selectivity, *α*^*t*^ an amplification factor and 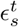 measurement noise. We performed a weighted least squares regression, assuming approximate independent variances 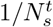 for the terms ln 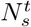, given that counts follow Poisson distributions [13]. We also estimated error bars for the selectivities *θ*_*s*_ by standard linear regression formulae [13].

To mitigate the effect of low counts, we add a pseudo-count of 1/2 to all counts before computing the empirical selectivities and before carrying out the inference [13].

### Structural contact predictions

The presence of a large epistatic effect between site positions is related to the three-dimensional proximity of the residues in the protein fold [20]. To quantify the strength of the epistatic effect we computed the difference between the fitness effect of double mutations and the sum of the effects of the two related single mutations, hence the expected additive fitness in absence epistasis. The genotype to fitness map in eq. (2), the fitness of a sequence is minus the energy *f* (*s*) = −*E*(*s*).

Considering a sequence *s*, the double mutant *v*_*ij*_ in position *i* and *j* and the single mutant *v*_*i*_ (and *v*_*j*_ resp.) in position *I* (and *j* resp.), we define the epistatic score as:

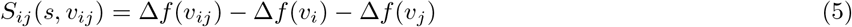

where Δ*f*(*v*_*ij*_) = *f*(*v*_*ij*_) − *f*(*s*) = − *E*(*v*_*ij*_) + *E*(*s*) = log(*P*(*v*_*ij*_)*/P*(*s*)) and similarly for Δ*f*(*v*_*i*_) and Δ*f*(*v*_*j*_). We substitute in *S*_*ij*_ and we average over each sequence *s* in the dataset and for each possible double mutants, obtaining:

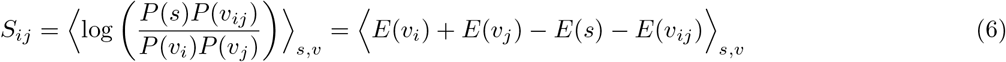

### Datasets

To assess the inferred genotype-phenotype map we used five datasets of mutational scan studies [2] that assess experimentally the mutational landscape of three different proteins. In the Olson et al. dataset [42], the effects of all single and double mutations between all positions in the IgG-binding domain of protein G (GB1) are quantified.

In this study, a library of all possible single and double amino-acid substitutions of the 55 sites of the GB1 protein domain is screened in a single round for the binding to an immunoglobulin fragment (IgG-Fc). The same protein and target pair were investigated by [43], who selected four positions in GB1 and exhaustively randomized them. Two more datasets come from [31] and [32], where a WW domain was randomized and selected for binding against its cognate peptide.

In the two studies, the wild type protein and the target are the same and, interestingly, the initial randomized libraries in the two datasets have about half of the sequences in common.

Finally, in [44], four positions of a variable antibody region are fully randomized and selected for binding against one of two targets: polyvinylpyrrolidone (PVP) or a short DNA loop with three cycles of selection.

The four positions are embedded into one of 23 possible frameworks. The main characteristics of the biological system and of the experimental settings are summarized in table 1. While in Olson *et al.* [42] the high mutant coverage (500) allows us to obtain a low sampling noise, the main limitations are due to the covered sequence space, limited to a maximum Hamming distance of two from the wild type sequence. In Boyer *et al.* [44] the fraction of covered sequence space is significant (16% of all possible sequences) and there are multiple rounds of selection but the obvious limitation comes from the small length of the mutated part of the sequence (4 a.a).

Compared to the previous datasets, the Fowler *et al.* and Araya *et al.* datasets have intermediate features where the covered sequence space is wider than in Olson et al. (average distance 3.4 and 4.3) and the number of selection rounds are respectively 3 and 4, but the sequencing depth is lower showing greater sampling noise.

## ACKNOWLEDGMENTS

The Authors acknowledge financial support from Marie Skƚodowska-Curie, grant agreement No. 734439 (INFERNET), and the Centro de Inmunologia Molecular of Cuba, and the Deparment of Physics of University of Havana, for their kind hospitality. We also thank C. Nizak and O. Rivoire for many interesting discussions.

## AUTHORS CONTRIBUTION

All authors contributed equally.

