## Supplementary Information for "Unsupervised inference of protein fitness landscape from deep mutational scan"

#### MATHEMATICAL APPENDIX

In this section we present the details of a more general model regarding the evolution of selection rounds in a deep mutational scan experiment, as well as the derivation and assumptions made to obtain the model discussed in the main-text.

Let sequence $s$ have selection probability $p_s$. If at round $t$ there are $N_s^t$ phages displaying sequence $s$, the probability that $n_s^t$ of them are selected for the next round follows a binomial law:

$$P(n_s^t|N_s^t, p_s) = \binom{N_s^t}{n_s^t} p_s^{n_s^t}(1-p_s)^{N_s^t - n_s^t} \tag{S.1}$$

The pool of selected phages is then amplified, so as to recover the same total number of phages of the original population. We assume that all sequences are amplified equally (*i.e.*, ignoring possible amplification biases). We use a bold symbol like $\mathbf{N}^t$ to denote the vector of all sequences at round $t$. Denoting $N_{\text{tot}} = \sum_s N_s^t$ and $n_{\text{tot}}^t = \sum_s n_s^t$, we can model this step with a multinomial law:

$$P(\mathbf{N}^{t+1}|\mathbf{n}^t) = \frac{N_{\text{tot}}!}{\prod_s N_s^{t+1}!} \prod_s \left(\frac{n_s^t}{n_{\text{tot}}^t}\right)^{N_s^{t+1}} \tag{S.2}$$

provided that $\sum_s N_s^{t+1} = N_{\text{tot}}$. Next, the probability of a time series of rounds $t = 0, 1, \dots, T$, is given by:

$$P(\mathbf{N}, \mathbf{n}|\mathbf{p}, \mathbf{N}^0) = \prod_{t=0}^{T-1} \sum_{\mathbf{n}^t} P(\mathbf{N}^{t+1}|\mathbf{n}^t) \prod_s P(n_s^t|N_s^t, p_s) \tag{S.3}$$

where bold symbols like $\mathbf{N}$ without a time super-script denote the set of all the values $N_s^t$ for all sequences and all rounds. If we have multiple replicates, we simply take a product of the above expression for all replicates.

The selection probabilities $p_s$ can be modeled assuming that each protein can be in one of two thermodynamic states: bound to the target or unbound. This is a simplification since it ignores additional thermodynamic states such as folded or unfolded [1]. In a two-state model, probabilities follow the Fermi-Dirac form:

$$p_s = \frac{1}{1 + \mathrm{e}^{E_s - \mu}} \tag{S.4}$$

where $\mu$ is the chemical potential (related to the logarithm of the target's concentration) and $E_s$ the binding energy. As mentioned in the maintext, we consider a two-body interaction expression for the dependence of $E_s$ upon the sequence:

$$E_s = -\sum_i h_i(s_i) - \sum_{i<j} J_{ij}(s_i, s_j) \tag{S.5}$$

The model is trained by maximizing the likelihood (S.3) with respect to the parameters $h_i(a)$ and $J_{ij}(a, b)$ in (S.5). However the model as defined so far remains intractable due to its dependence on $\mathbf{n}$ (which must be marginalized because it is not observed). We thus adopt a *deterministic binding* approximation, and assume that $n_s^t = p_s N_s^t$. Taking logarithms in equation (S.3), employing Stirling's approximation, and omitting terms independent of $p_s$, we obtain the log-likelihood:

$$\mathcal{L} = \ln P(\mathbf{N}|\mathbf{p}, \mathbf{N}^0) = \sum_{s,t} \mathcal{L}_s^t \tag{S.6}$$

where

$$\mathcal{L}_s^t = N_s^{t+1} \ln \frac{N_s^t p_s}{\sum_\sigma N_\sigma^t p_\sigma} \tag{S.7}$$

Notice that $\mathcal{L}$ is not concave with respect to the parameters. Therefore inferring the model by maximum likelihood can be challenging since the optimization can get stuck at local maxima. However, in the *rare binding* regime, we have that $\mu - E_s$ is large and negative so that $p_s \ll 1$ for most sequences. This is consistent with the observation that empirically a very small fraction of the phage population is selected in each round. In this case, we can approximate $p_s \approx \mathrm{e}^{\mu - E_s}$. With this approximation, we obtain

$$\mathcal{L}_s^t = N_s^{t+1} \ln \frac{N_s^t \mathrm{e}^{-E_s}}{\sum_\sigma N_\sigma^t \mathrm{e}^{-E_\sigma}} \tag{S.8}$$

Observe that: 1) under this approximation, $\mathcal{L}$ is a concave function of the parameters; 2) $\mu$ simplifies out. This way we obtained the model discussed in the paper.

### Mutual Information

One of the most commonly used non-parametric measures of statistical dependency between two residues is the mutual information between the distributions of amino acids occurring in the two positions [2], reflects the extent to which knowledge of the amino acid at one position allows us to predict the identity of the amino acid at the other position.

Let $f_i(a)$ and $f_{ij}(a,b)$ be the selectivity-weighted frequencies to observe amino acid $a$ at position $i$ $f_i(a)$ and the co-frequencies to have amino acid $a$, $b$ at position $i$ and $j$ respectively:

$$f_i(a) = \frac{1}{M} \sum_s \theta_s \delta_{s_i,a}, \quad f_{ij}(a,b) = \frac{1}{M} \sum_s \theta_s \delta_{s_i,a} \delta_{s_j,b} \tag{S.9}$$

The Mutual Information between postion i and j is defined as :

$$\mathrm{MI}_{ij} = -\sum_{a,b} f_{ij}(a,b) \log \left( \frac{f_{ij}(a,b)}{f_i(a) f_j(b)} \right) \tag{S.10}$$

### SUPPLEMENTARY FIGURES

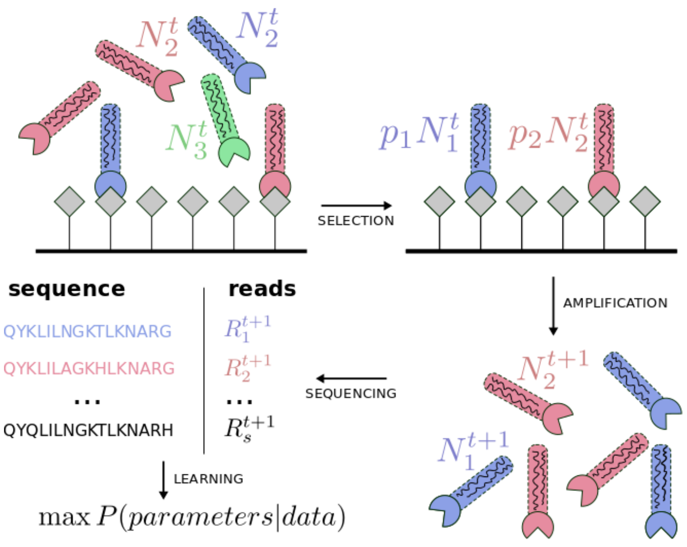


FIG. S1. **Experimental pipeline and modelization scheme.** Phages displaying sequence variant $s$ are present with abundances $N_s^t$ at round $t$. They are exposed to a binding target for which the displayed proteins exhibit some affinity. The variants that bind are selected (with a sequence dependent probability $p_s$), while those that do not bind are washed away. The selected variants are amplified so that the next round can start from a total abundance similar to the previous round. A small sub-sample of the amplified phages are collected for sequencing. The reads taken at each round are then used to train a model, as explained in the text.

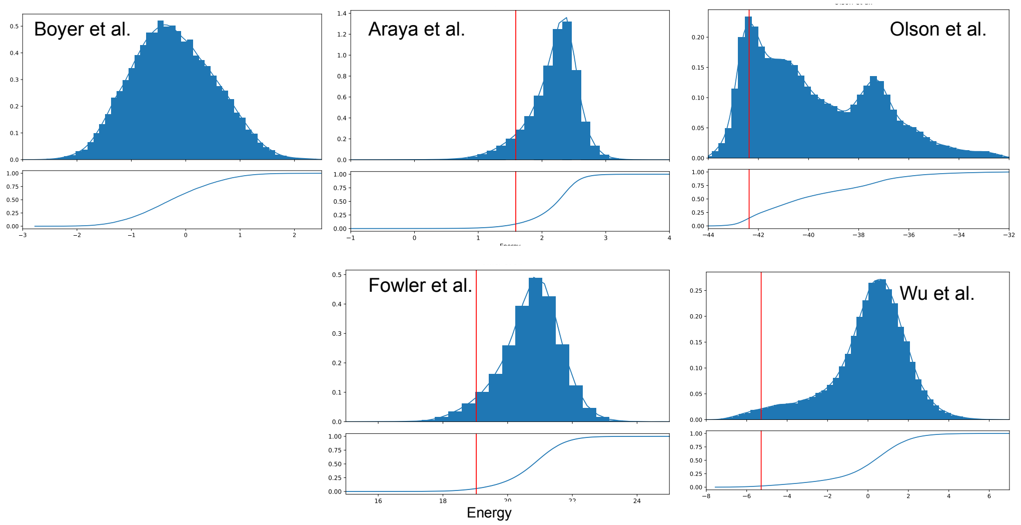


FIG. S2. **Distribution of mutational effects.** For each dataset is plotted the histogram of the energy (inferred fitness) distrbutions (panel above) and the cumulative distribution (panel below), the red line show the value of the wild type (except for the Boyer et al. dataset).

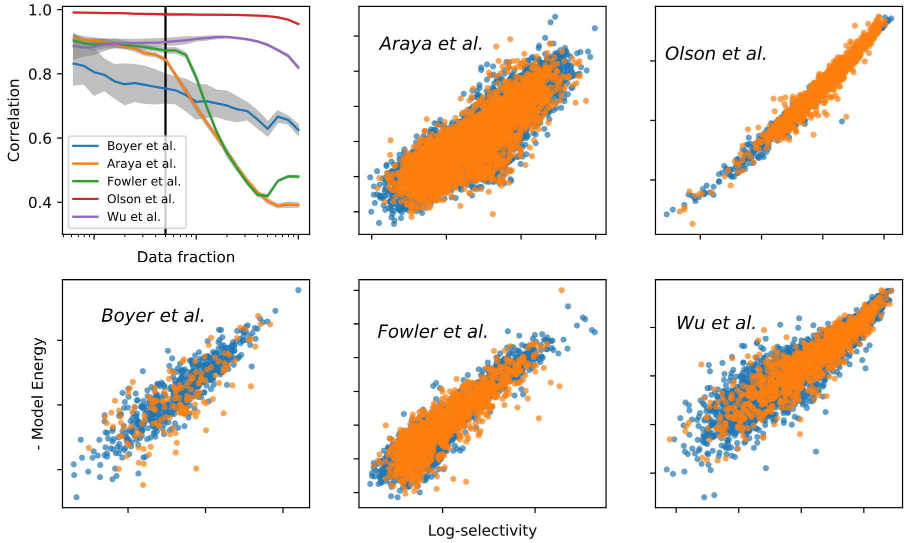


FIG. S3. **Binding energy and selectivity.** Scatter plots of the inferred energy vs the log-selectivity (computed from the sequencing reads), in blue the in-sample set and in orange the out-of-sample sequences in each dataset. In the first panel are shown the Pearson correlations between energy and the log-selectivity for different filtering thresholds on sequence errors. On the x-axis the fraction of the sequences used to compute the correlation, the vertical line shows the fraction of tested sequences used in the scatter plots shown in the other panels ($f = 0.05$). The Pearson correlations coefficients for the out-of-sample sequences are in clockwise order 0.84, 0.98, 0.90, 0.88, 0.72.

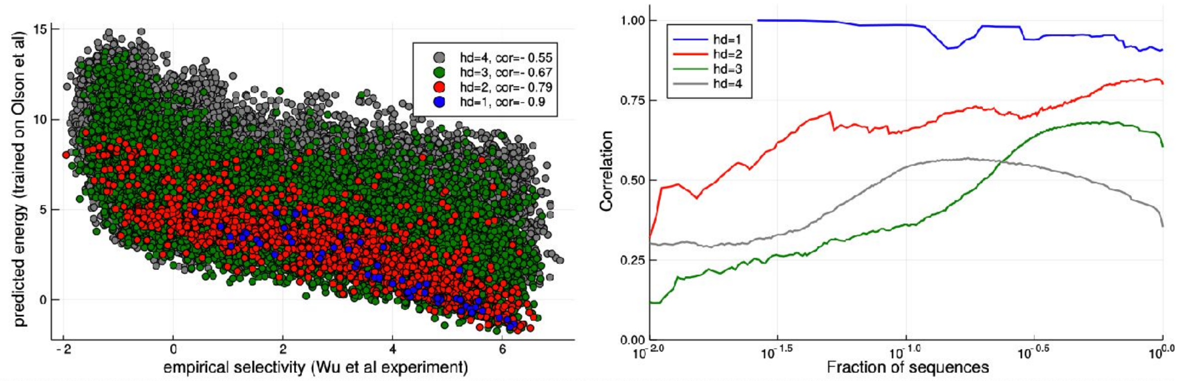


FIG. S4. **Model trained in Olson et al. dataset and tested on the Wu et al. dataset**. In Olson et al. [3] and the Wu et al. [4], the IgG-binding domain of protein G (GB1) is selected for binding to immunoglobulin G fragment crystallizable (IgG-Fc). Olson et al. generated an exhaustive library containing all 1-point and 2-point mutants away from the wild-type while Wu et al. exhaustively varied four chosen positions of the GB1 protein and performed 2 rounds of selection.**a)** The $x$-axis are the empirical selectivities of GB1 variants measured in the Wu *et al.* experiment ($\theta_{\text{Wu}}$). The $y$-axis are the energies of these variants as predicted by a model trained on the Olson *et al.* dataset ($E_{\text{Olson}}$). **b)** Pearson correlation coefficient between predicted energies $E_{\text{Olson}}$ and empirical log-selectivity $\theta_{\text{Wu}}$ for different filtering thresholds of sampling error. The color legend distinguishes variants by their Hamming distance to the wild-type GB1 sequence. The Pearson correlation ($\rho$) is highest for sequences with Hamming distances of 1 ($\rho = 0.9$) and 2 ($\rho = 0.79$) to the wild type, since these are the variants covered in the Olson et al. dataset used for training. The model still achieves good predictions for sequences with more mutations, though with lower Pearson correlations of $\rho = 0.67$ for Hamming distance 3 and $\rho = 0.55$ for Hamming distance 4.

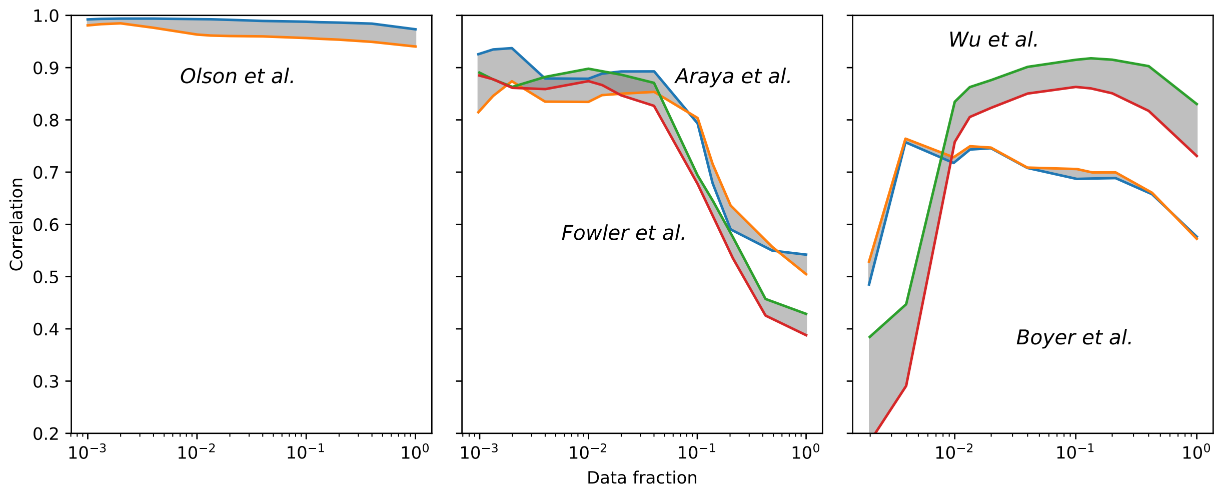


FIG. S5. **Global Epistasis model [5].** The comparison of the non-epistatic model and the Global Epistatic model outcome on the five datasets. For each dataset is shown the correlation of the Global Epistatic model (upper line) and the linear non-epistatic site model (lower line) and the gap between the two is highlighted in gray. To compare with the specific epistatic model, see the panels (a-c) of the figure (3) of the main text. From left to right the broadness of the screened library increases and the length of the mutated sequence reduces (see table I). Broader the library, more distant mutants from the wild type are screened. The Global Epistasis has a reduced accuracy in the selectivity predictions as broadness of the library increases.

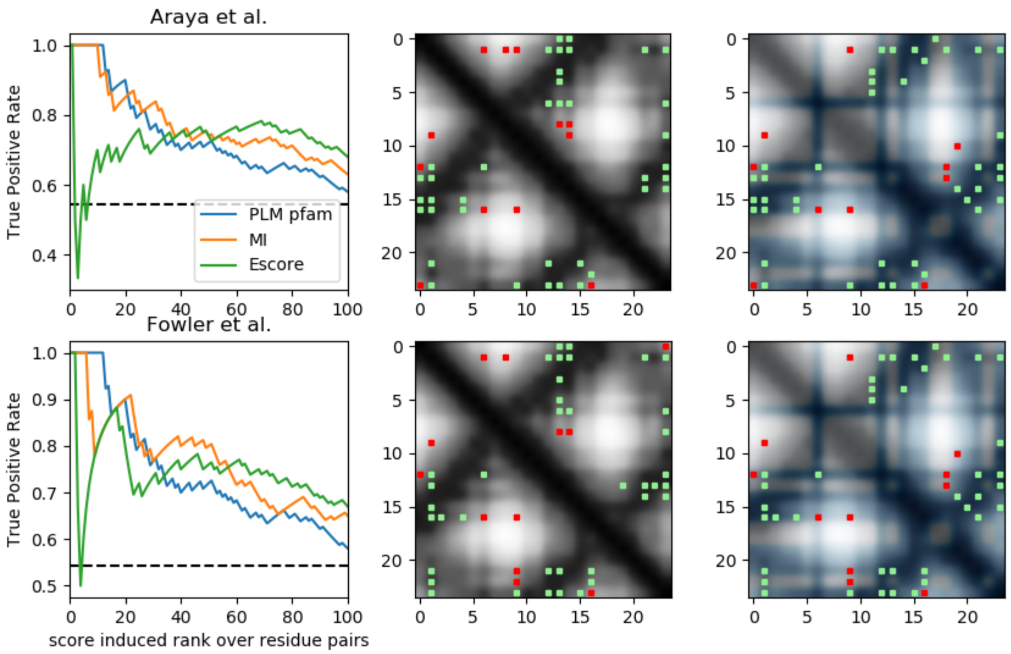


FIG. S6. **Contacts predictions of WW domain.** The figure shows the structure contacts predictions of the WW domain from the Araya et al. (upper panels) and Fowler et al. (lower panels) DMS experiment (tested on the crystal structure PDB id: 1jmq). Panels on the left: the ROC curves of the predicted contacts with the epistatic score computed from the inferred model and the weighted Mutual Information (see methods section), is shown also the prediction of Direct Coupling Analysis from homologous sequences of the WW domain family (using the PlmDCA algorithm [6]). Interestingly the accuracies of the predictions from the DMS experiments are close to the standard DCA ones and are very similar to each other. Other Panels: contact map of the first predicted contacts. In gray-scale is displayed the distances between residue heavy atoms, the green dots are the true positives (heavy atoms distance less than 8Å) and the reds are false positives. Central panels show in the upper triangular part the Mutual Information predictions while in the lower triangular part the Epistatic score. Right panels show in the upper triangular part the homology plmDCA predictions while in the lower triangular part the Epistatic score, in blue are highlighted the residues on the binding surface (less than 3Å to the Fc domain of human IgG).
